# Metabolomic Profiling of Serum Biomarkers in Women with Polycystic Ovary Syndrome: Insights from an Untargeted Approach

**DOI:** 10.64898/2026.02.26.707762

**Authors:** Jalpa Patel, Hiral Chaudhary, Sonal Panchal, Rushikesh Joshi

## Abstract

**Background:** Polycystic ovary syndrome (PCOS) is a complex endocrine disorder characterized by metabolic dysregulation. Identifying serum biomarkers can enhance our understanding of its pathophysiology. This study employs an untargeted metabolomic approach to investigate metabolic alterations in PCOS.

**Methods:** Serum samples were collected from 71 women with PCOS and 54 healthy controls. Untargeted Metabolomic profiling was performed using liquid chromatography-mass spectrometry (LC-MS) to identify differentially abundant metabolites. Pathway analysis was conducted to identify key metabolic disruptions, and correlations between identified metabolites and clinical parameters were assessed.

**Results:** The metabolomics analysis identified 24 upregulated and 17 downregulated metabolites in PCOS compared with controls. These metabolites mainly include glycerophospholipids, fatty acids, sphingolipids, peptides, ceramides, and steroids. Pathway analysis indicated that these metabolites were enriched in pathways including bile acid biosynthesis, glycerolipid metabolism, tryptophan metabolism, the citric acid cycle, and fatty acid metabolism. Increased levels of branched-chain and aromatic amino acids suggested potential links to insulin resistance. Disruptions in bile acid metabolism pointed to altered gut microbiome interactions. Additionally, metabolites related to oxidative stress and mitochondrial function indicated metabolic dysfunction. Correlation analyses revealed associations between altered metabolites and clinical markers such as insulin resistance and androgen levels.

**Conclusion:** This study reveals distinct serum metabolic alterations in PCOS, emphasizing their association with insulin resistance and inflammation. These findings highlight the potential of metabolomics to identify novel biomarkers for early diagnosis and to develop targeted therapeutic strategies.

## 1. Introduction

Polycystic Ovary Syndrome (PCOS), a common endocrine disorder affecting 5–10% of reproductive-aged women globally, poses significant reproductive and metabolic health challenges (1)(2). Characterised by hyperandrogenism, irregular menstrual cycles, and polycystic ovaries, PCOS is also associated with metabolic disturbances such as insulin resistance, obesity, and dyslipidaemia, increasing the risk of type 2 diabetes and cardiovascular diseases (3)(4). Although PCOS is highly prevalent, its causes and molecular mechanisms are not yet fully understood, emphasizing the urgent need for improved diagnostic and therapeutic approaches.

Traditional diagnostic approaches for PCOS, mainly centered on hormonal and clinical indicators, frequently overlook the comprehensive metabolic disturbances associated with the disorder (5). Given that insulin resistance is a core feature of PCOS, a shift towards metabolomics, the study of small molecules resulting from cellular processes, is crucial (6). Untargeted metabolomics provides a comprehensive platform for analyzing a wide range of metabolites, facilitating the understanding of metabolic changes and the discovery of new biomarkers (7)(8). Mass spectrometry is essential for untargeted metabolomics, allowing accurate profiling of serum samples to identify metabolic signatures linked to PCOS.

Previous research has highlighted disruptions in metabolic pathways involving lipid and amino acid metabolism, as well as energy production, in individuals with PCOS (9)(10). However, many of these studies have used targeted metabolomic analyses, limiting the identification of more extensive metabolic abnormalities that contribute to the phenotypic diversity of PCOS (11). To address these knowledge gaps, this study employs an untargeted metabolomic approach to enhance the understanding of metabolic heterogeneity in PCOS.

This research identifies metabolic changes in women with PCOS versus healthy controls and explores how these relate to clinical features, aiming to find early diagnostic markers or treatment targets. While some global studies exist, few are from India, and ethnicity influences metabolic profiles, meaning single-centre results may not reflect broader diversity. This study seeks to fill these gaps, deepen understanding of PCOS’s molecular basis, and support development of diagnostic tools for early detection and intervention, reducing long-term health risks.

## 2. Methodology

### 2.1 Subjects and Ethical Approval

This study included 125 women, categorized into two groups: 71 with PCOS and 54 healthy controls. The participants, aged 20 to 42 years, were recruited individually from Dr. Nagori’s Institute and the Health Centre at Gujarat University in Ahmedabad, India, between December 2019 and March 2023. To maintain consistency in diagnosing PCOS, we aligned our criteria with the internationally recognised Rotterdam Criteria, specifically adapted for Indian women. Participants in the PCOS group were required to show at least two of the following symptoms: irregular or anovulation, clinical or biochemical evidence of hyperandrogenism, and the presence of polycystic ovaries, defined as having at least 12 follicles between 2–9 mm or an ovarian volume exceeding 10 cm³. The control group consisted of women without any previous PCOS diagnosis, allowing for a distinct separation between the groups.

The Ethical Committee at Gujarat University approved this research ethically (Reference: GU IEC(NIV)/02/Ph.D./007). All processes were carried out in strict alignment with applicable guidelines and ethical standards. Participants had to provide informed consent to join; all study details were communicated verbally in both English and Gujarati, and written consent was collected through a bilingual consent form prior to their involvement in the study activities.

### 2.2 Sample Collection and Biochemical Assessment

Blood specimens were collected from participants during the follicular phase (days 2-5) of their menstrual cycle after an overnight fast. This timing was intentional, designed to enhance conditions for biochemical evaluations and serum metabolomic analysis. Serum samples for the research were collected between January 2023 and December 2024, representing metabolic profiles from both PCOS and healthy controls throughout this period and informing them of the study’s purpose. After collection, the specimens were centrifuged at 2500 rpm for 15 minutes, and the serum was stored at -80°C until analysis. Biochemical investigations included measuring luteinizing hormone (LH), follicle-stimulating hormone (FSH), and 17β-estradiol, along with thyroid-stimulating hormone (TSH) and prolactin (PRL) to rule out thyroid and prolactin disorders. Testosterone (T) and dehydroepiandrosterone sulfate (DHEA-S) levels were also measured to evaluate hyperandrogenemia, using chemiluminescent assays at an accredited clinical laboratory.

### 2.3 Chemicals

All chemicals met analytical-reagent-grade standards. Ammonium acetate was sourced from Sigma. Acetonitrile, water, and formic acid of LC-MS grade were obtained from J.T. Baker.

### 2.4 Sample Preparation

Our analytical approach was based on the protocol established by Patel et al. (2025)(12), which details sample preparation and mass spectrometric conditions for the study. Briefly, to prepare serum samples, transfer 100 μL of serum to a microcentrifuge tube. Add 400 μL of chilled acetonitrile to precipitate proteins and extract metabolites. Vortex vigorously to mix thoroughly, then incubate at -20°C for 15 minutes to enhance protein precipitation. Centrifuge at 12,000 rpm and 4°C for 10 minutes to separate the supernatant containing extracted metabolites from the pellet. Transfer the clear supernatant to a new microcentrifuge tube without disturbing the pellet. Vacuum-dry the supernatant to remove acetonitrile and concentrate the metabolites. Resuspend in 300 μL of acetonitrile and filter through a 0.22 μm nylon syringe filter to ensure sample purity before LC-MS analysis.

A quality control (QC) sample was prepared by combining equal volumes (10 μL each) of serum from all study samples to assess system stability and method performance. This QC sample is analysed after every five serum samples, helping to confirm data reliability and reduce potential false positives in untargeted metabolomics on the LC-MS platform. Although untargeted analysis provides broad metabolite coverage, it sometimes lacks precise quantitative data and can yield false positives without reference standards. A targeted analysis of specific metabolites could deliver more accurate insights into group variations. This robust validation process ensures the accuracy, reproducibility, and reliability of serum metabolomic analysis, supporting LC-MS analysis and a detailed exploration of the metabolomic landscape.

### 2.5 Instrumentation

The analysis employed a Bruker Elute UHPLC system with Hystar 5.0 SR1 software and a Bruker Daltonics AmaZon Speed™ mass spectrometer, with data processed via Data Analysis 5.2, all from Bruker Daltonics. Metabolites were separated using a Waters UPLC ACQUITY BEH C18 column (1.7 μm, 50×2.1 mm) with a guard. Mobile phases differed for positive and negative modes: positive used Water with 0.1% formic acid (A) and acetonitrile with 0.1% formic acid (B); negative used Water with 0.1% ammonium acetate (A) and acetonitrile with 0.1% ammonium acetate (B). Samples were at 30 °C, with 5 μL injected and separated by a gradient from 5% B to 15%, then back, over 32 minutes at 300 μL/min, at 30 °C. Mass detection in both modes used Auto MS(n), with m/z 50–1500, capillary at 4.5 kV, nebulizer at 29.0 psi, dry gas at 10.0 L/min, and temperature 126.9 °C, ensuring precise metabolomic profiling.

### 2.6 Metabolomic Data Analysis

The MSMS spectra were analyzed using respected libraries, including HMDB, Metabolomics Workbench, and MoNA, ensuring accurate metabolite identification. Data processing used MetaboAnalyst 6.0, involving logarithmic transformation, auto-scaling, and sum normalization. The refined data was then analyzed with 2D and 3D PCA, clearly distinguishing PCOS and control groups.

After the unsupervised analysis, a focused examination using Orthogonal Partial Least Squares Discriminant Analysis (OPLS-DA) provided a deeper investigation of metabolic differences between the two groups. The quality of OPLS-DA models was assessed using goodness-of-fit (R2Y) and predictive capability (Q2). VIP plots identified metabolites significantly contributing to group separation, with VIP values above 1.0 considered potential biomarkers. Permutation tests with 2000 repetitions ensured that the observed OPLS-DA separation was not coincidental (p < 0.05). The statistical significance of metabolite differences was assessed using paired parametric t-tests and the Mann–Whitney test, with the Bonferroni correction to address multiple testing. Results with P values and false discovery rates (FDR; q-values) below 0.05 were deemed statistically significant, validating the study’s conclusions.

## 3. RESULTS

### 3.1 Biochemical and clinical profiles of participants

The study compared demographic and biochemical parameters between women with PCOS (n=71) and control subjects (n=54) (Table 1). Women with PCOS were significantly older (28.54 ± 4.35 years vs. 24.58 ± 4.92 years, p < 0.0001) and had a higher BMI (25.97 ± 5.13 kg/m² vs. 22.74 ± 4.58 kg/m², p < 0.001). The PCOS group had higher LH levels (7.17 ± 4.49 mIU/ml vs. 5.02 ± 4.25 mIU/ml, p = 0.008). The groups did not differ significantly in testosterone, estradiol, FSH, PRL, LH/FSH ratio, DHEA-S, and TSH levels. Additional parameters for the PCOS group included fasting glucose (84.82 ± 13.88 mg/dl), postprandial glucose (113.47 ± 28.21 mg/dl), fasting insulin (9.98 ± 3.91 µIU/ml), postprandial insulin (92.11 ± 68.69 µIU/ml), and HOMA-IR (2.05 ± 0.85), which were not measured in controls.

**Table 1.**
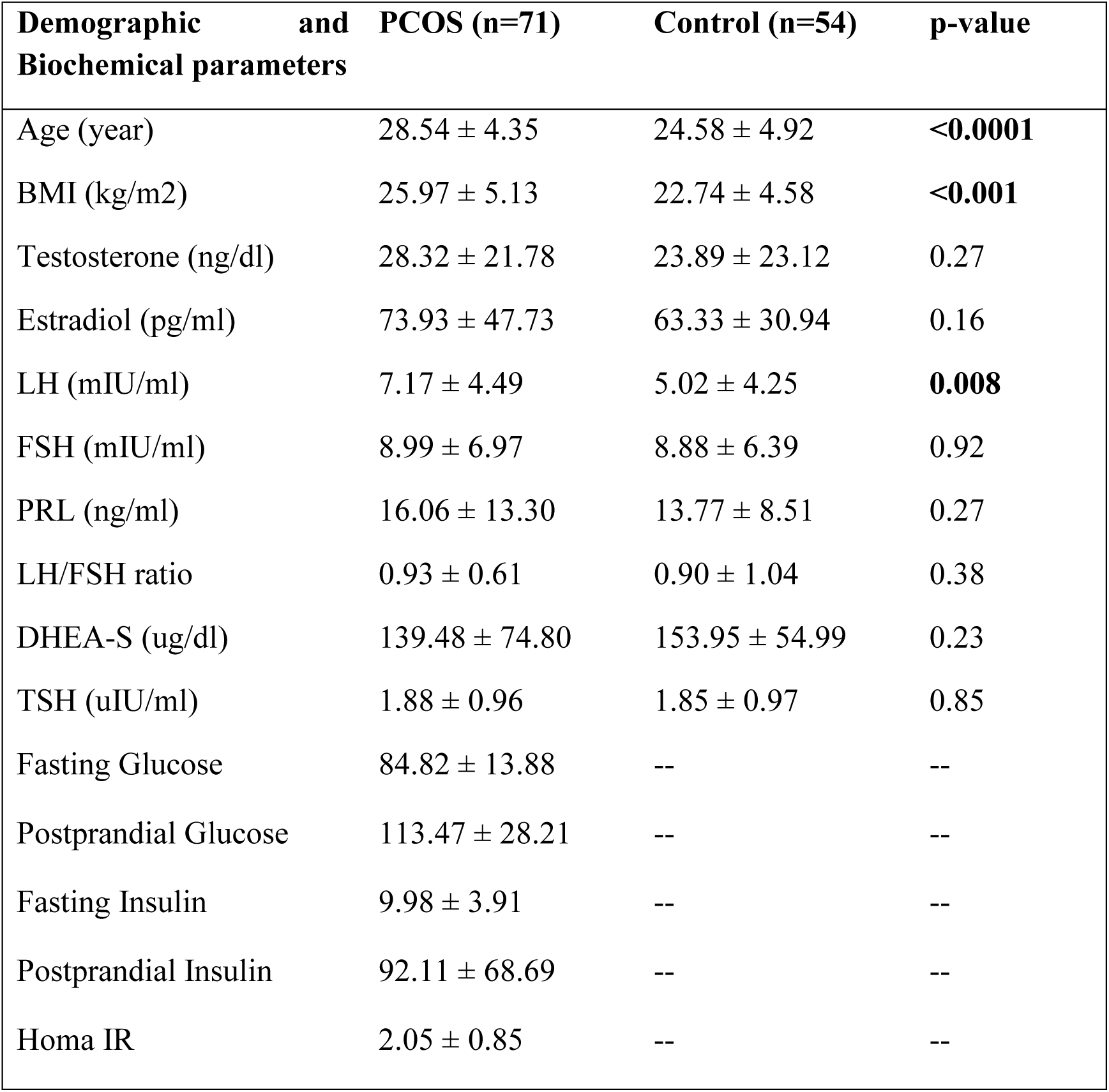
Demographic and biochemical parameters of study participants for serum metabolomic study

| Demographic and Biochemical parameters | PCOS (n=71) | Control (n=54) | p-value |
| --- | --- | --- | --- |
| Age (year) | $28.54 \pm 4.35$ | $24.58 \pm 4.92$ | **<0.0001** |
| BMI (kg/m<sup>2</sup>) | $25.97 \pm 5.13$ | $22.74 \pm 4.58$ | **<0.001** |
| Testosterone (ng/dl) | $28.32 \pm 21.78$ | $23.89 \pm 23.12$ | 0.27 |
| Estradiol (pg/ml) | $73.93 \pm 47.73$ | $63.33 \pm 30.94$ | 0.16 |
| LH (mIU/ml) | $7.17 \pm 4.49$ | $5.02 \pm 4.25$ | **0.008** |
| FSH (mIU/ml) | $8.99 \pm 6.97$ | $8.88 \pm 6.39$ | 0.92 |
| PRL (ng/ml) | $16.06 \pm 13.30$ | $13.77 \pm 8.51$ | 0.27 |
| LH/FSH ratio | $0.93 \pm 0.61$ | $0.90 \pm 1.04$ | 0.38 |
| DHEA-S (ug/dl) | $139.48 \pm 74.80$ | $153.95 \pm 54.99$ | 0.23 |
| TSH (uIU/ml) | $1.88 \pm 0.96$ | $1.85 \pm 0.97$ | 0.85 |
| Fasting Glucose | $84.82 \pm 13.88$ | -- | -- |
| Postprandial Glucose | $113.47 \pm 28.21$ | -- | -- |
| Fasting Insulin | $9.98 \pm 3.91$ | -- | -- |
| Postprandial Insulin | $92.11 \pm 68.69$ | -- | -- |
| Homa IR | $2.05 \pm 0.85$ | -- | -- |

### 3.2 Differential Serum Metabolite Expression in PCOS women

After identifying metabolites from databases, we found sets of metabolites in both groups. The Venn diagram (Figure 1(a)) shows 61 metabolites unique to PCOS, 58 unique to controls, and 21 shared metabolites, highlighting distinct metabolic profiles and shared metabolites. PCA analysis (Figure 1(b)) reveals clear separation between groups, with PC1 (49.6%) and PC2 (6.2%) accounting for most of the variation. Samples cluster separately, indicating significant metabolic differences that may serve as biomarker.

**Figure 1.**
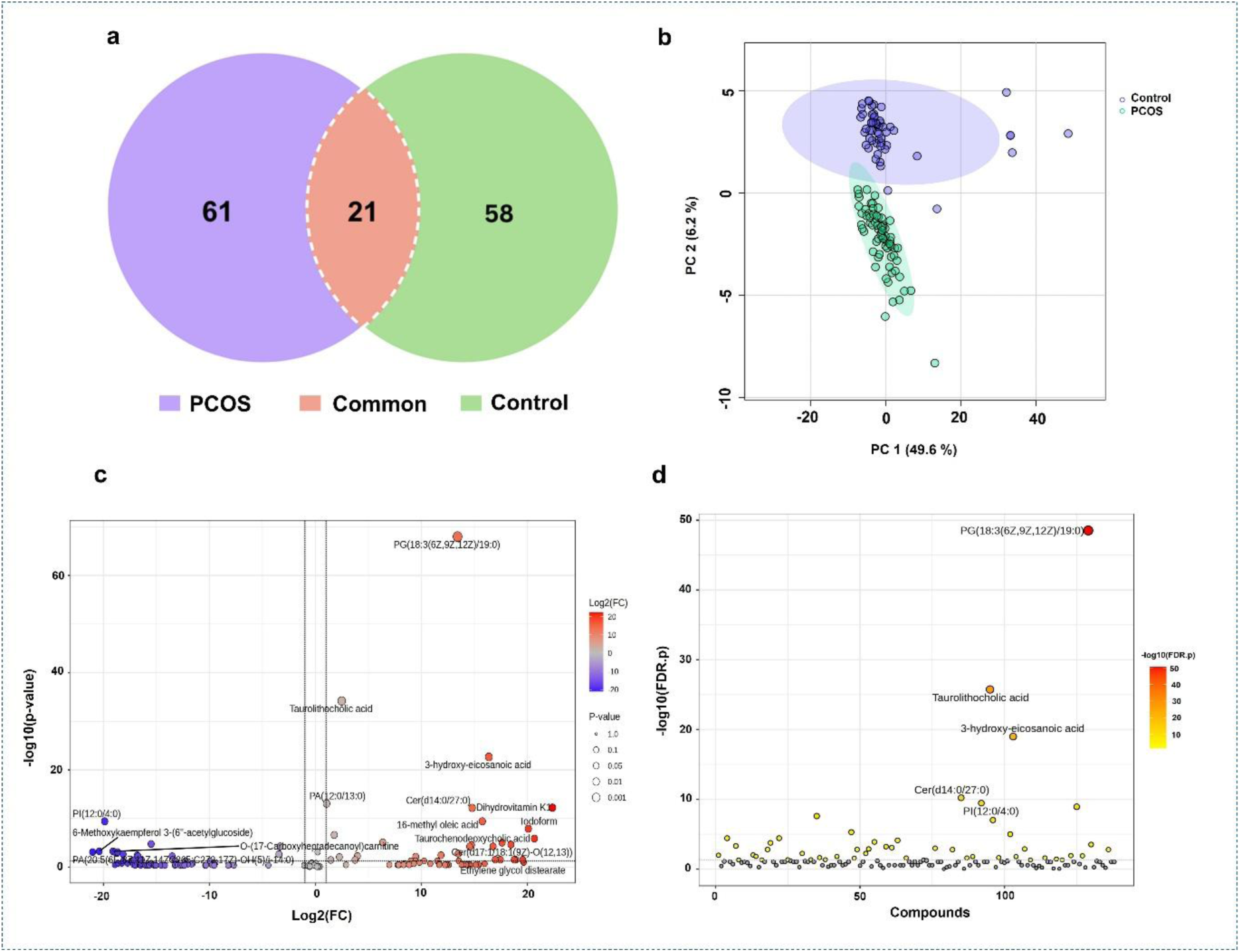
Metabolomic profiling of serum samples in PCOS and control groups shows 61 metabolites unique to PCOS, 58 unique to controls, and 21 common, indicating distinct and overlapping profiles. PCA plot separates PCOS (cyan) and controls (purple), with PC1 explaining 49.6% of variance and PC2 6.2%, highlighting clear group differences. A volcano plot of -log10(p-value) versus log2(fold change) highlights metabolites like PG(18:3(6Z,9Z,12Z)/19:0), Taurolithocholic acid, and 3-hydroxy-eicosanoic acid with significant fold changes and low p-values. A t-test with -log10(FDR-adjusted p-value) emphasises significant metabolites, suggesting potential PCOS biomarkers.

Supplementary Table 1 presents the identified metabolites, detailing m/z, class, ionisation mode, fold change (FC), log2(FC), and regulation in control versus PCOS groups, thereby highlighting metabolic alterations associated with PCOS. These metabolites are categorized into classes such as glycerophospholipids, ceramides, sphingolipids, bile acids, among others, reflecting diverse affected pathways. The ionisation mode indicates detection conditions and chemical properties.

Notably, 3-hydroxy-eicosanoic acid and Asn-Tyr are highly upregulated in PCOS, with FC values of 60.98 and 41.31, respectively. Conversely, PI (12:0/4:0) and 3’-deoxy-cytidine-5’-triphosphate are significantly downregulated, with FC values of 0.009 and 0.093. Fold change illustrates the level differences: FC > 1 indicates upregulation, while FC < 1 signifies downregulation (represented by red for up, green for down). For example, 3-hydroxy-eicosanoic acid (FC=60.98, log2(FC)=5.93) and Asn-Tyr (FC=41.31, log2(FC)=5.369) are upregulated, whereas PI (12:0/4:0) (FC=0.009, log2(FC)=-6.8) and 3’-deoxy-cytidine-5’-triphosphate (FC=0.093, log2(FC)=-3.43) are downregulated, indicating metabolic pathway disruptions.

The study revealed significant differences in metabolite levels between groups. A volcano plot illustrates these changes, with log2 fold change on the x-axis and -log10 p-value on the y-axis (Figure 1(c)). Several metabolites, including phosphatidylglycerol PG(18:3(6Z,9Z,12Z)/19:0), showed marked downregulation in PCOS. Additionally, taurolithocholic acid, 3-hydroxy-eicosanoic acid, Cer(d14:0/27:0), dihydro vitamin K1, 16-methyl oleic acid, iodoform, and tauro chenodeoxycholic acid demonstrated moderate decreases.

Conversely, metabolites such as PI(12:0/4:0), 3’-deoxy-cytidine-5’-triphosphate, and 6-methoxy kaempferol 3-O-6’’-acetylglucoside were upregulated, suggesting potential biomarkers or therapeutic targets. T-test analysis (Figure 1(d)) confirms significant differences between control and PCOS groups, with PG(18:3(6Z,9Z,12Z)/19:0) among the most significant, along with taurolithocholic acid, 3-hydroxy-eicosanoic acid, Cer(d14:0/27:0), and PI(12:0/4:0), as indicated by color gradients reflecting the magnitude of change.

Figure 2 shows the top 20 metabolites as violin plots. Metabolites like PG (18:3(6Z,9Z,12Z)/19:0), Taurolithocholic acid, and 3-hydroxy-eicosanoic acid are lower in PCOS. Cer(d14:0/27:0), Dihydrovitamin K1, and 16-methyl oleic acid are also decreased. In contrast, PI (12:0/4:0), PA (12:0/13:0), and 3’-Deoxy-cytidine-5’-triphosphate are higher in PCOS. Other metabolites include Iodoform, FAD, Cer(d17:1/16:0), and Tauro chenodeoxycholic acid. Additionally, PA (20:5(6E,8Z,11Z,14Z,17Z)/C26:5,172)-OH (5)/1:14:0, 6-Methoxykaempferol 3-(6’-acetylglucoside), and Phosphotidylinositol-3,4,5-triphosphate show distinct patterns. These differences suggest altered metabolism in PCOS, indicating potential biomarkers and pathways. All metabolites had significant p-values (<0.01), confirming robustness and their data shown in the supplementary table 2 PLS-DA analysis of serum profiles showed clear separation between groups, with 33.6% and 22.1% of the variance explained, as indicated by the confidence ellipses (Figure 3(a)). OPLS-DA further refined this, isolating classification-related variation and confirming group distinctions (Figure 3(b)).

**Figure 2.**
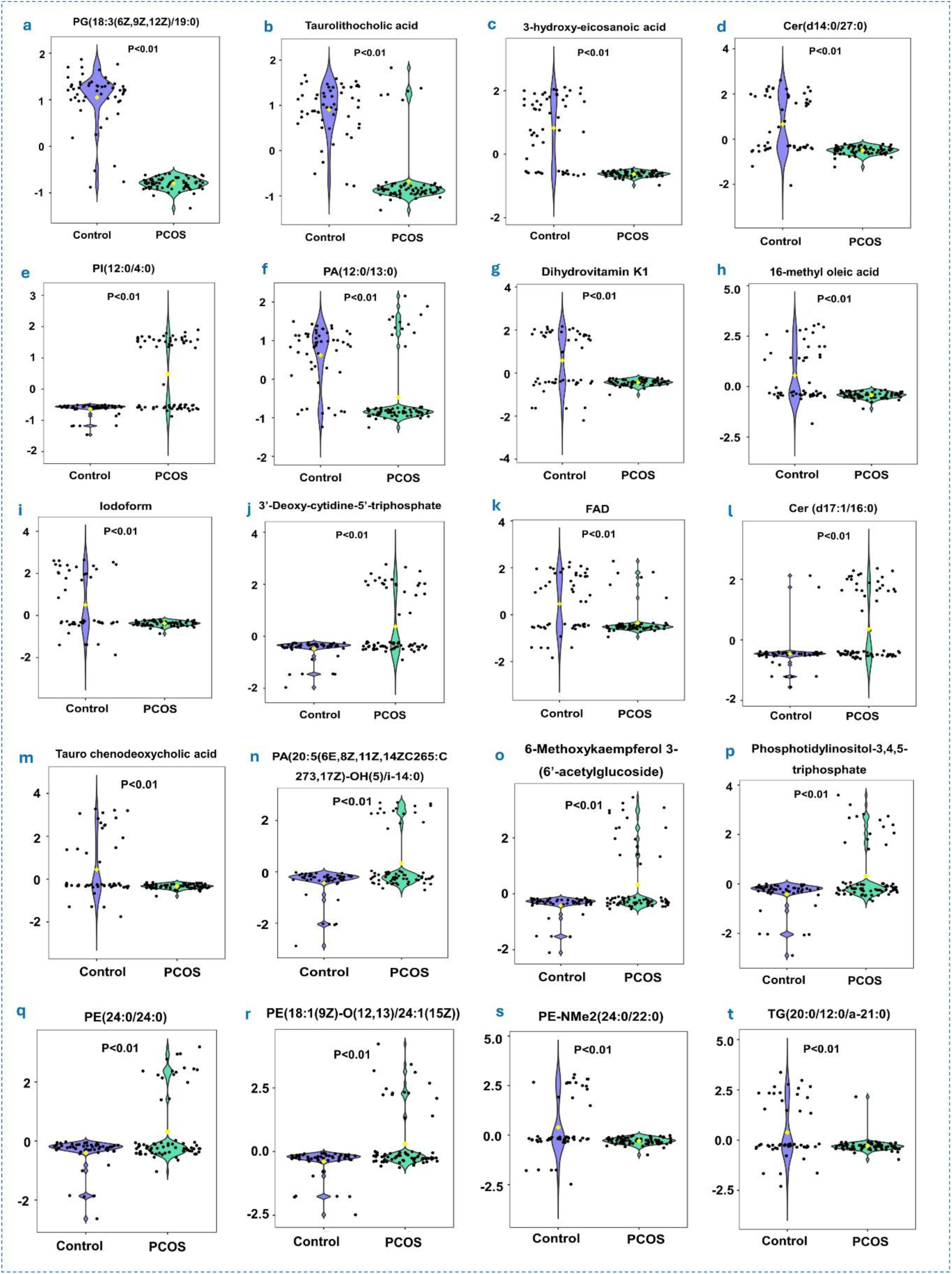
Violin plots of the top 20 differentially expressed serum metabolites between control and PCOS groups. Panels (a-t) display violin plots for the top 20 metabolites, showing the distribution and expression levels in control versus PCOS samples. The metabolites include PG(18:3(6Z,9Z,12Z)/19:0) (a), Taurolithocholic acid (b), 3-hydroxy-eicosanoic acid (c), Cer(d14:0/27:0) (d), PI(12:0/4:0) (e), PA(12:0/13:0) (f), Dihydrovitamin K1 (g), 16-methyl oleic acid (h), Iodoform (i), 3’-Deoxy-cytidine-5’-triphosphate (j), FAD (k), Cer(d17:1/16:0) (l), Taurochenodeoxycholic acid (m), PA(20:5(6E,8Z,11Z,14Z)/C26:5(17Z)-OH(5)/14:0) (n), 6-Methoxykaempferol 3-(6’-acetyl glucoside) (o), Phosphatidylinositol-3,4,5-triphosphate (p), PE(24:0/24:0) (q), PE(18:1(9Z)/O-12:13(4Z)/1(15Z)) (r), PE-NMe2(24:0/22:0) (s), and TG(20:0/12:0/a-21:0) (t). Each plot includes a p-value indicating significant differences (P<0.01) between the groups, highlighting the potential of these metabolites as biomarkers for PCOS.

**Figure 3.**
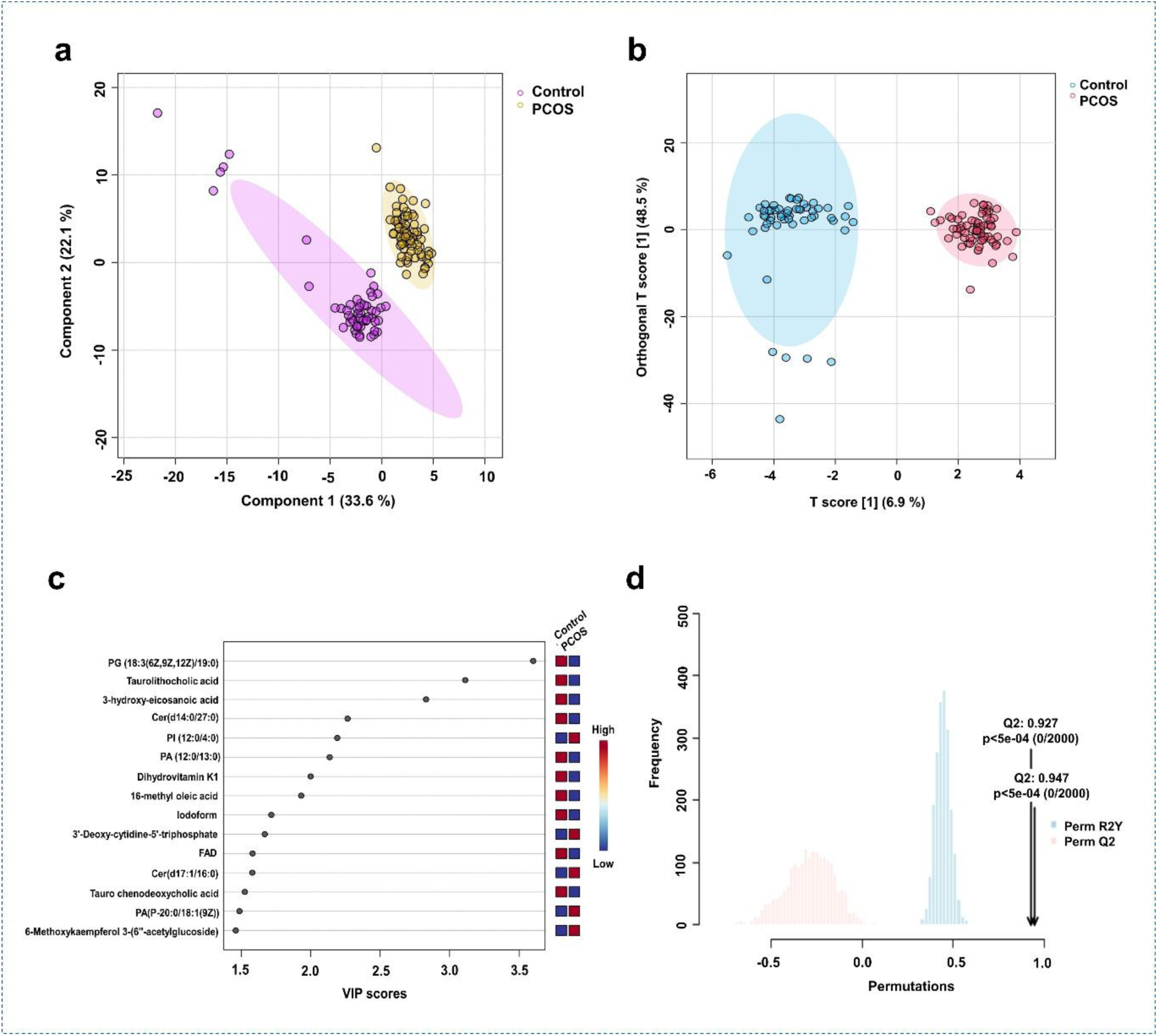
Serum Untargeted Metabolomic Profiling. (a) PLS-DA score plot separates control (purple) and PCOS (yellow) groups by the first two components, explaining 33.6% and 22.1% of variance. Ellipses show 95% confidence. (b) OPLS-DA plot separates control (blue) and PCOS (red) groups along the T scores, with orthogonal T scores capturing non-classification variation. Ellipses indicate 95% confidence. (c) VIP scores highlight key metabolites, with higher scores indicating greater importance. The heatmap shows relative metabolite levels in control (blue) and PCOS (red). (d) Permutation tests validate PLS-DA, with R2Y=0.947 and Q2=0.927 from 2000 permutations, exceeding chance (p < 5e-04).

Key metabolites for separation were identified using VIP scores (Figure 3(c)). High VIP scores for metabolites such as PG (18:3(6Z,9Z,12Z)/19:0), Taurolithocholic acid, and 3-hydroxy-eicosanoic acid highlighted their importance in distinguishing control from PCOS groups. A heatmap displayed their relative abundance, clearly distinguishing the control (blue) and PCOS (red) groups. Permutation testing validated the robustness of the PLS-DA model (Figure 3(d)). R2Y and Q2 values from 2000 permutations showed the observed R2Y (0.947) and Q2 (0.927) were significantly higher than chance (p < 5e-04), confirming the model’s effectiveness in differentiating control and PCOS.

### 3.3 Feature Importance and Significance Analysis of Serum Metabolites

To investigate differences between control and PCOS groups, we conducted feature importance and significance analyses. The feature importance plot (Figure 4(a)) illustrated the correlation between primary component scores, p[1], and their correlation coefficients, p(corr)[1], for the metabolites. This linear relationship, shown by magenta circles, indicated consistent feature importance across components, highlighting the reliability of these features in distinguishing control from PCOS groups.

**Figure 4.**
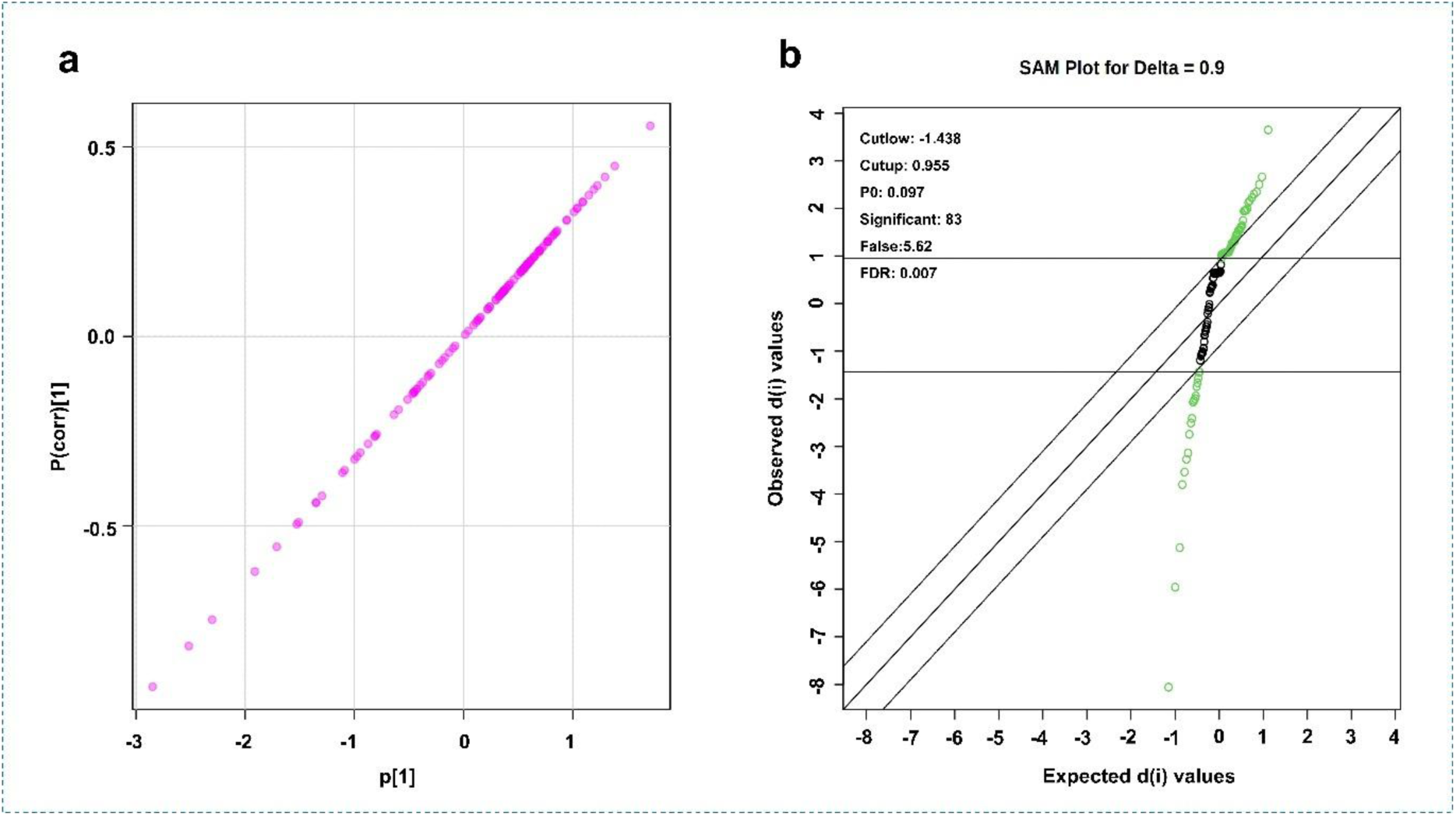
Feature Importance and Significance Analysis in Control and PCOS Groups. (a) Feature importance plot shows the correlation between component scores, p[1], and correlation coefficients, p(corr)[1], for metabolites. The magenta circles indicate consistent feature importance across components. (b) SAM plot for delta = 0.9 compares observed d(i) values to expected under no difference. Features outside cutoff values of -1.438 and 0.955, marked in green and black, are significant. The analysis finds 83 significant features with an FDR of 0.007.

The SAM plot (Figure 4(b)) confirmed these findings by identifying features differentiating the two groups. With a delta of 0.9, it plotted observed d(i) against expected values under no difference, identifying 83 significant features with cutoffs at -1.438 and 0.955. The false discovery rate was 0.007, with 5.62 false positives and a feature proportion of 0.097. Features above the upper cutoff (green circles) and below the lower cutoff (black circles) were significant, distinguishing control and PCOS groups. Metabolites that differentiate these groups were found, with high feature importance and significant SAM features revealing metabolic differences related to PCOS. These results highlight metabolomic profiling’s role in identifying PCOS biomarkers and pathways, and in validating methods for clinical and research use.

### 3.4 Serum Biomarker Analysis of PCOS

The study found that predictive accuracy increased with the number of features, plateauing around 30-50 features (Figure 5(a)). Receiving operative Curve (ROC) curves for 5, 10, 15, 50, and 100 features showed AUC from 0.878 to 0.999, indicating strong sensitivity and specificity, with more features generally improving performance.

**Figure 5.**
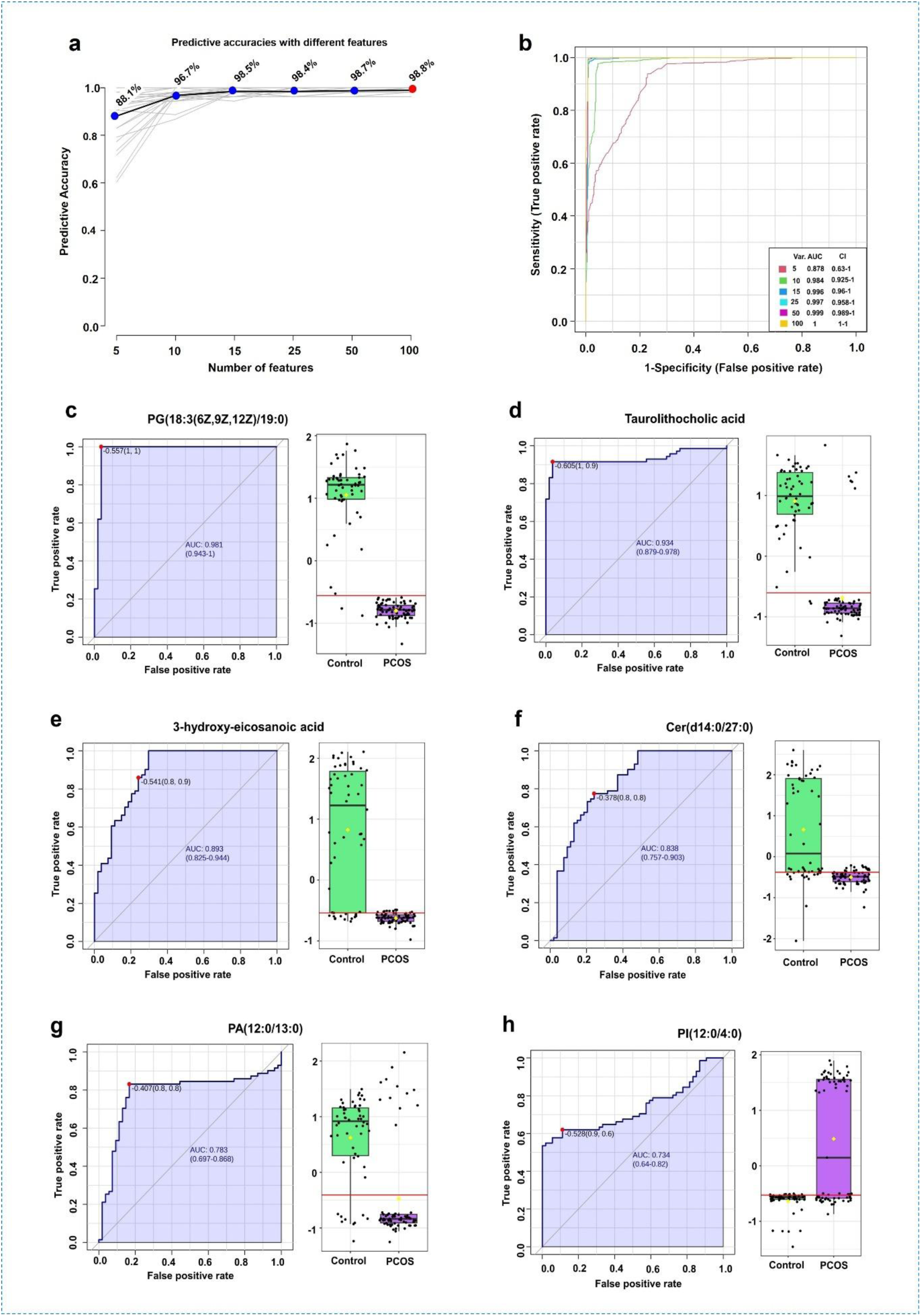
Serum biomarker analysis. Panel (a) shows the predictive accuracy as a function of the number of features, indicating a plateau in accuracy improvement around 30-50 features. Panel (b) presents ROC curves for varying features (5, 10, 15, 50, and 100), with AUC values ranging from 0.878 to 0.999, demonstrating high sensitivity and specificity. Panels (c-h) illustrate the ROC curves and box plots for individual metabolites, including PG(18:3(6Z,9Z,12Z)/19:0), Taurolithocholic acid, 3-hydroxy-eicosanoic acid, Cer(d14:0/27:0), PA(12:0/13:0), and PI(12:0/4:0), highlighting their varying degrees of effectiveness in distinguishing between control and PCOS samples.

Metabolite analysis identified several with notable discriminatory power, such as PG (18:3(6Z,9Z,12Z)/19:0) (AUC=0.981) and taurolithocholic acid (AUC=0.934). Other metabolites like 3-hydroxy-eicosanoic acid (AUC=0.893) and Cer(d14:0/27:0) (AUC=0.838) also showed good discrimination. Moderate markers included PA (12:0/13:0) and PI (12:0/4:0) with AUCs of 0.783 and 0.734. These metabolites could serve as PCOS biomarkers, aiding early and accurate detection.

### 3.5 Correlational analysis between serum metabolites and clinical parameters

The heatmap (Figure 6) shows a correlation analysis of serum metabolites in women with PCOS and controls, including BMI, Estradiol, Testosterone, DHEAS, TSH, LH, FSH, PRL, and LH/FSH ratio. Columns represent samples; rows depict metabolites, with colors indicating physiological and hormonal parameters. Intensity shows concentrations; dark green for high BMI, dark blue for high Estradiol. Hierarchical clustering groups similar metabolites, highlighting affected pathways. Comparing PCOS to controls reveals higher cholesterol derivatives and ceramides in higher-BMI samples, indicating a metabolic signature linked to body mass. Estradiol correlates with 4-hydroxy-17β-estradiol and Ceramides, revealing estrogen differences. Testosterone links to metabolites like 6-chlorotryptophan, relevant in hyperandrogenism. DHEAS associates with DG (22:5) and DG (20:5), showing metabolic modulation. TSH influences PE (24:0/24:0); LH and FSH affect O-(17-Carboxyheptadecanoyl) carnitine, important for reproductive and metabolic functions, with group differences. PRL and LH/FSH ratio impact Ethylene glycol distearate and NAD, indicating prolactin pathways. Variations in Diacylglycerols and Ceramides relate to lipid metabolism influenced by hormones. Steroid metabolites and vitamins like Dihydrovitamin K1 and FAD reflect hormonal and physiological states.

**Figure 6.**
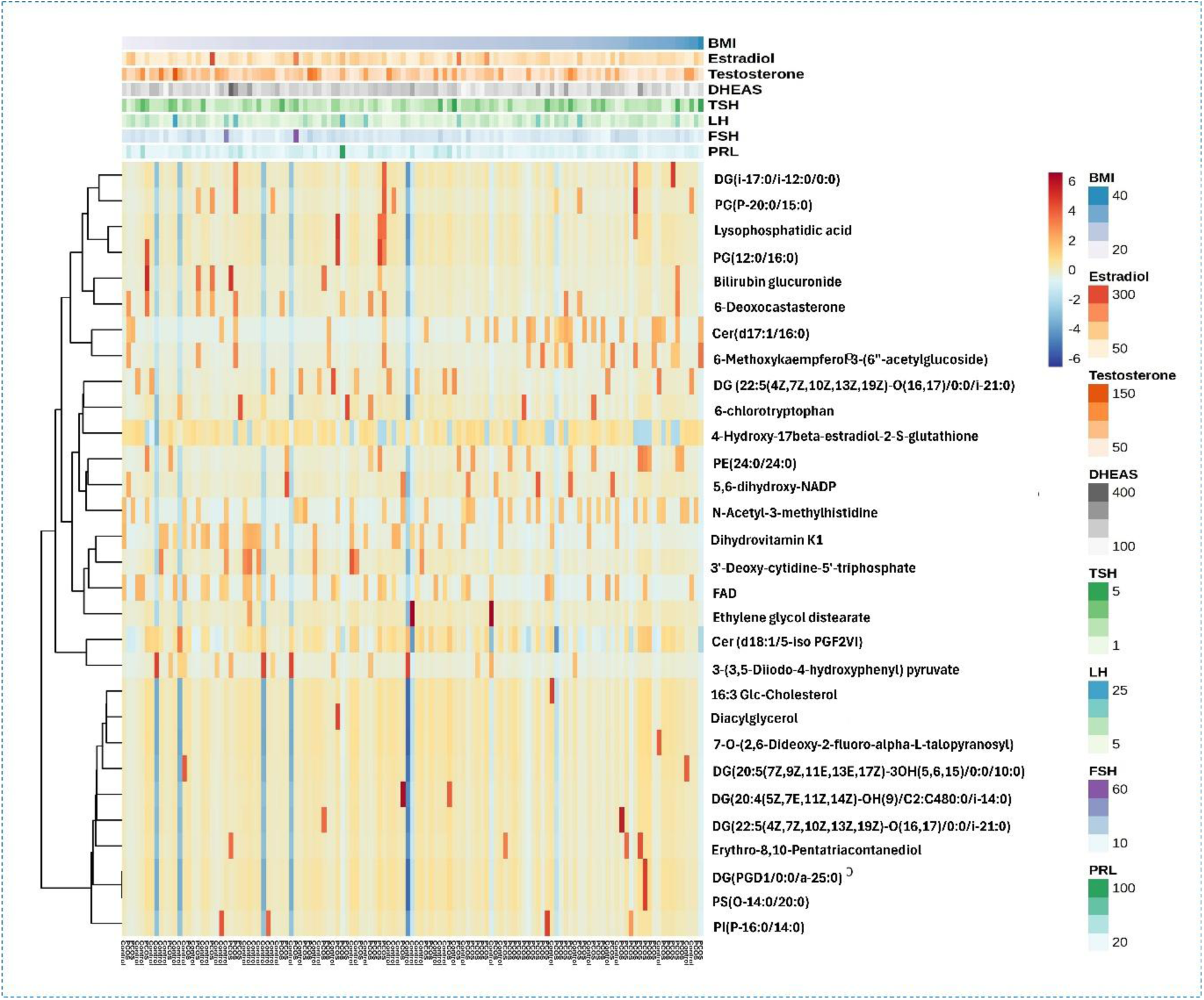
Heatmap of hierarchical cluster analysis of serum metabolites linked to demographic and hormonal variables in women with PCOS and controls. Columns are samples, rows are metabolites. Top annotations show parameters (BMI, Estradiol, Testosterone, DHEAS, TSH, LH, FSH, PRL, LH/FSH), with colours indicating levels. The heatmap colour scale ranges from red (high) to blue (low). Clustering uncovers metabolic profiles and biomarkers associated with PCOS, highlighting hormonal correlations.

### 3.6 Pathway enrichment analysis of serum metabolomics of PCOS

Figure 7 shows the top 25 enriched metabolic pathways comparing control and PCOS groups, with controls against PCOS. The x-axis shows the enrichment ratio, indicating pathway overrepresentation in PCOS versus controls. The color gradient from yellow to red indicates p-value significance, with darker shades for lower p-values.

**Figure 7.**
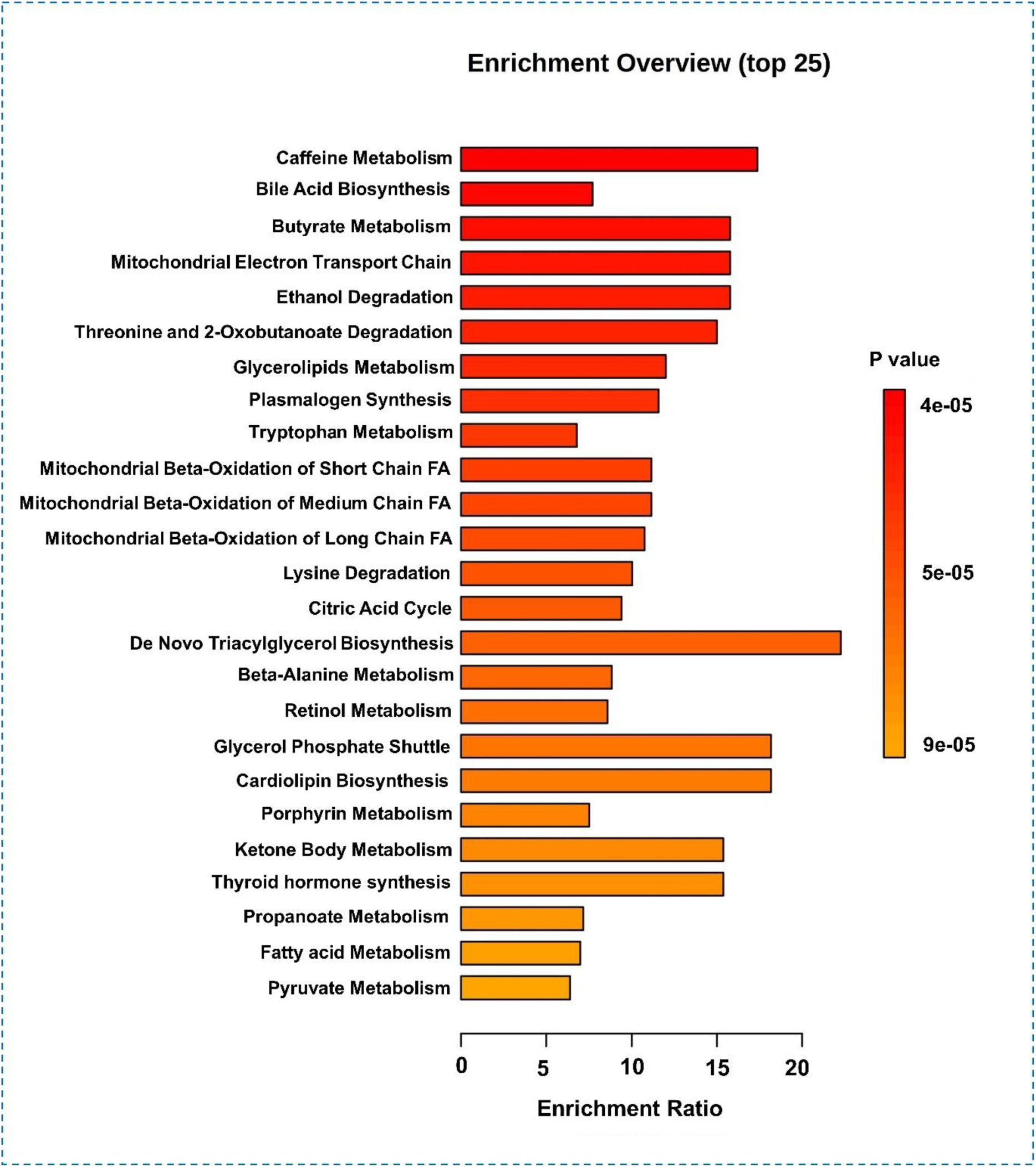
Pathway enrichment analysis of the top 25 metabolic pathways in control and PCOS groups shows enrichment ratios and pathways in a bar plot. Yellow to red indicates p-value significance, with darker being more significant. Key pathways like Caffeine Metabolism, Bile Acid Biosynthesis, and Butyrate Metabolism have high enrichment ratios and low p-values, highlighting metabolic differences. The analysis highlights alterations in mitochondrial function, lipid metabolism, and amino acid degradation in PCOS.

Key findings reveal notable metabolic changes in PCOS, including high enrichment in caffeine metabolism, bile acid and butyrate biosynthesis, and significant disruptions in amino acid, lipid, and energy pathways. Mitochondrial dysfunction and altered alcohol metabolism are evident, highlighting potential biomarkers and therapeutic targets.

## 4. DISCUSSION

PCOS is a common endocrine disorder affecting 8-13% of women of reproductive age, with symptoms like menstrual irregularities, hyperandrogenism, and polycystic ovaries (13). The prevalence of disorders and manifestations varies significantly across geographical locations, influenced by distinct environmental, lifestyle, and dietary factors (14)(15). Given the complex interplay of genetic, metabolic, and environmental factors in PCOS, this study used untargeted metabolomics to examine biochemical differences between healthy controls and women with PCOS.

The serum metabolomic study shows biochemical and clinical differences between women with PCOS and controls, deepening the understanding of PCOS-related metabolic dysregulation. The PCOS group had higher BMI and LH levels, consistent with known hormonal imbalances and increased BMI in PCOS (16)(17). Elevated LH levels in PCOS corroborate findings by *Malini et. al.,* 2028 (18), underscoring the endocrine abnormalities typical in PCOS.

Multivariate analysis revealed significant metabolic changes in PCOS, with glycerophospholipids, fatty acyls, and dipeptides notably upregulated. Increased levels of 3-hydroxy-eicosanoic acid and Asn-Tyr support previous findings of disrupted fatty acid metabolism (19). Conversely, the downregulation of PI (12:0/4:0) and 3’-Deoxy-cytidine-5’-triphosphate is consistent with the observed alterations in lipid metabolism among PCOS patients (20).

In our study, women with PCOS had significantly lower taurolithocholic acid (TLCA) levels than healthy controls, indicating altered bile acid metabolism linked to the condition. This reduction may relate to metabolic issues in PCOS, like insulin resistance and dyslipidemia, as bile acids regulate lipid and glucose metabolism (21)(22). Bile acids like TLCA have anti-inflammatory properties, and reduced levels could worsen the chronic inflammation and oxidative stress seen in PCOS (23). The gut microbiota, which modulates bile acid composition, might be implicated, as dysbiosis is common in PCOS and could further disrupt bile acid homeostasis (24). Bile acids are recognized as signalling molecules interacting with nuclear receptors like FXR to regulate glucose and lipid metabolism.

Further research into bile acid dysregulation in PCOS is needed, and targeting these pathways could lead to new therapies for this syndrome.

Our study found lower levels of taurolithocholic acid and other metabolites, including sphingolipids (e.g., Cer), fatty acids (3-hydroxy-eicosanoic acid), and glycerophospholipids (PG), in women with PCOS compared to controls. These results support the literature showing disrupted lipid metabolism in PCOS. Sphingolipids, especially ceramides, are vital for cell signaling and apoptosis, and their dysregulation links to metabolic issues like insulin resistance, common in PCOS (25)(26). The observed reduction in ceramide levels suggests impaired sphingolipid metabolism, which may exacerbate insulin resistance and metabolic dysfunction in PCOS (27).

Furthermore, fatty acyls like 3-hydroxy-eicosanoic acid are vital for energy balance and inflammation. Decreased levels may signal issues in fatty acid oxidation and inflammation, key factors in PCOS pathophysiology (28). Glycerophospholipids are vital for cell membranes and metabolic signalling. The decreased PG (18:3(6Z,9Z,12Z)/19:0) levels in our study suggest changes in membrane lipids, potentially affecting metabolism and contributing to PCOS-related disturbances (29).

The current study found that certain metabolites, including glycerophospholipids such as PI (12:0/4:0) and PA (12:0/13:0), metabolites from the pentose phosphate pathway such as 3’-deoxy cytidine-5’-triphosphate, and fatty acyls such as o-(17-carboxyheptadecanoyl) carnitine, were higher and upregulated in the PCOS group compared to controls. This may reflect compensatory mechanisms or activation of alternative metabolic pathways in response to PCOS-related metabolic stress. Elevated glycerophospholipids may indicate increased membrane turnover or remodelling, possibly as a cellular response to maintain membrane integrity and function under metabolic stress (10). The increased pentose phosphate pathway metabolites indicate a higher demand for nucleotide synthesis and redox balance, likely due to oxidative stress (30). Similarly, higher levels of fatty acyls such as o-(17-carboxyheptadecanoyl) carnitine could indicate altered fatty acid oxidation and mitochondrial function in PCOS (31). These contrasting findings highlight the complex metabolic alterations in PCOS and underscore the need for comprehensive metabolic profiling to fully understand its pathophysiology. This aligns with previous studies which reported aberrant lipid metabolism in systemic conditions, and emphasised the role of oxidative stress and redox balance in metabolic disorders (32)(33). Our results contribute to the growing body of evidence that PCOS involves multifaceted metabolic disturbances and point towards potential therapeutic targets in lipid and bile acid metabolism for managing PCOS.

Pathway enrichment analysis revealed significant changes in metabolic pathways between control and PCOS groups. Notably, caffeine metabolism, bile acid biosynthesis, and butyrate metabolism were enriched in the PCOS group, indicating disrupted lipid and energy metabolism. This aligns with a previous study, which reported mitochondrial dysfunction and altered lipid metabolism in PCOS (34). Enrichment of mitochondrial beta-oxidation pathways and the citric acid cycle emphasizes the disturbances in energy production and utilization in PCOS, supporting the hypothesis of widespread metabolic disruptions in this condition (35). The enrichment of these pathways underscores the importance of these metabolic alterations in the pathophysiology of PCOS.

This study’s strength lies in its novel contribution to metabolomics in PCOS among the Indian population, offering region-specific insights that improve understanding of the disorder’s metabolic basis. Variations in ethnicity and environment influence metabolic profiles, making our findings valuable to global PCOS knowledge. The metabolic changes highlight the need for region-specific studies, as genetic, dietary, and lifestyle factors influence metabolomic results. However, the cross-sectional design limits causal conclusions, and a small sample size may restrict broad applicability. Future research with larger groups, longitudinal data, and gut microbiome analysis is essential to better understand metabolic and microbial interactions in PCOS.

Despite limitations, our study advances understanding of PCOS metabolism and highlights potential biomarkers. The metabolic signatures suggest avenues for targeted interventions and precision medicine, improving diagnostics and treatment strategies. Future research should validate these biomarkers and explore their use in early detection, prognosis, and monitoring response in PCOS patients.

## Abbreviations

PCOS: Polycystic ovary syndrome
BMI: body mass index
FSH: follicle stimulating hormone
LH: luteinizing hormone
TSH: thyroid stimulating hormone
DHEAS: dehydroepiandrosterone sulfate
PRL: prolactin
E2: estradiol
T: testosterone
LC-MS: Liquid chromatography-mass spectrometry
HMDB: Human metabolome database
QC: Quality control
OPLS-DA: Orthogonal Partial Least Squares Discriminant Analysis
PLS-DA: Partial Least Squares Discriminant Analysis
PCA: Principal component analysis
FDR: False discovery rate
FC: Fold change
SAM: Significance Analysis of Microarrays
VIP: Variable importance in projection
ROC: Receiver Operating Characteristic
AUC: Area under curve

## Acknowledgement

Jalpa Patel is grateful to the Department of Education, Government of Gujarat, India for providing Scheme of Developing High-Quality Research (SHODH) fellowship. Hiral Chaudhary is thankful to Ministry of Education, Government of India for providing CSIR-UGC-NET fellowship. The authors are thankful to the Department of Biochemistry and Forensic Science, Gujarat University for laboratory facilities.

## Funding

This research was funded by the Anusandhan National Research Foundation of India (grant number ANRF/PAIR/2025/000008).

## Disclosure statement

The authors declare that they have no competing interests.

## Author contribution

JP and HC assisted with data collection, writing, and manuscript preparation. SP assisted in sample collection. RJ carried out the critical review.

## Data Availability

The mass spectrometry data described in the manuscript is accessible at https://doi.org/10.6084/m9.figshare.28938701.v1.

## Notes

https://doi.org/10.6084/m9.figshare.28938701.v1.

### Competing Interest Statement

The authors have declared no competing interest.

## References

1. Rao P, Bhide P. Controversies in the diagnosis of polycystic ovary syndrome. Ther Adv Reprod Heal. 2020;14:263349412091303.

2. Fahs D, Salloum D, Nasrallah M, Ghazeeri G. Polycystic Ovary Syndrome: Pathophysiology and Controversies in Diagnosis. Diagnostics. 2023;13(9):1–13.

3. Chaudhary H, Ph D, Patel J, Ph D, Jain NK, Ph D, et al. Impact of CYP19A1 genetic variations on polycystic ovary syndrome : fi ndings from a case-control study. Fertil Steril Sci [Internet]. 2025; Available from: 10.1016/j.xfss.2025.03.005

4. Patil M. The Prevalence of PCOS in South Asia. Fertil Reprod. 2023;05(04):190–190.

5. Nair GR, Jishamol K, Navami S. Advancements In Diagnostic Approaches For Polycystic Ovary Syndrome ( PCOS ): A Comprehensive Review. 2023;11(6):903–8.

6. Naigaonkar A, Dadachanji R, Kumari M, Mukherjee S. Insight into metabolic dysregulation of polycystic ovary syndrome utilizing metabolomic signatures: a narrative review. Crit Rev Clin Lab Sci [Internet]. 2024;62(2):85–112. Available from: 10.1080/10408363.2024.2430775

7. Wishart DS. Metabolomics for investigating physiological and pathophysiological processes. Physiol Rev. 2019;99(4):1819–75.

8. Aderemi AV, Ayeleso AO, Oyedapo OO, Mukwevho E. Metabolomics: A scoping review of its role as a tool for disease biomarker discovery in selected non-communicable diseases. Metabolites. 2021;11(7).

9. Zhao Y, Fu L, Li R, Wang LN, Yang Y, Liu NN, et al. Metabolic profiles characterizing different phenotypes of polycystic ovary syndrome: Plasma metabolomics analysis. BMC Med. 2012;10.

10. Yu Y, Tan P, Zhuang Z, Wang Z, Zhu L, Qiu R, et al. Untargeted metabolomic approach to study the serum metabolites in women with polycystic ovary syndrome. BMC Med Genomics [Internet]. 2021;14(1):1–15. Available from: 10.1186/s12920-021-01058-y

11. Chang AY, Lalia AZ, Jenkins GD, Dutta T, Carter RE, Singh RJ, et al. Combining a nontargeted and targeted metabolomics approach to identify metabolic pathways significantly altered in polycystic ovary syndrome [Internet]. Vol. 71, Metabolism: Clinical and Experimental. Elsevier Inc.; 2017. 52–63 p. Available from: 10.1016/j.metabol.2017.03.002

12. Joshi R, Patel J, Chaudhary H. Comparing the Metabolomic Landscape of Polycystic Ovary Syndrome within Urban and Rural Environments [Internet]. 2025. Available from: https://figshare.com/articles/dataset/_b_Comparing_the_Metabolomic_Landscape_of_Polycystic_Ovary_Syndrome_within_Urban_and_Rural_Environments_b_/28891628

13. Wolf WM, Wattick RA, Kinkade ON, Olfert MD. Geographical prevalence of polycystic ovary syndrome as determined by region and race/ethnicity. Int J Environ Res Public Health. 2018;15(11):1–13.

14. Patel J, Chaudhary H, Panchal S, Joshi R. Plasticizer exposure and reproductive dysfunction: Assessing bisphenol A and phthalate esters impact on ovarian reserve in women with PCOS-associated infertility. Reprod Toxicol [Internet]. 2025;135(February):108949. Available from: 10.1016/j.reprotox.2025.108949

15. Patel J, Chaudhary H, Panchal S, Joshi T, Joshi R. Endocrine-disrupting chemicals and hormonal profiles in PCOS women : A comparative study between urban and rural environment. Reprod Toxicol [Internet]. 2024;125(February):108562. Available from: 10.1016/j.reprotox.2024.108562

16. Chaudhary H, Patel J, Jain NK, Panchal S, Laddha N, Joshi R. Association of FTO gene variant rs9939609 with polycystic ovary syndrome from Gujarat, India. BMC Med Genomics. 2023;16(1):1–9.

17. Gene H, Chaudhary H, Patel J, Jain NK, Panchal S, Laddha N. Investigating the interplay between AMH gene polymorphism rs10407022 and clinical indicators in polycystic ovary syndrome. Hum Gene [Internet]. 2024;40(January):201279. Available from: 10.1016/j.humgen.2024.201279

18. Malini NA, Roy George K. Evaluation of different ranges of LH:FSH ratios in polycystic ovarian syndrome (PCOS) – Clinical based case control study. Gen Comp Endocrinol [Internet]. 2018;260:51–7. Available from: 10.1016/j.ygcen.2017.12.007

19. Cree-Green M, Carreau AM, Rahat H, Garcia-Reyes Y, Bergman BC, Pyle L, et al. Amino acid and fatty acid metabolomic profile during fasting and hyperinsulinemia in girls with polycystic ovarian syndrome. Am J Physiol - Endocrinol Metab. 2019;316(5):E707–18.

20. Chen YX, Zhang XJ, Huang J, Zhou SJ, Liu F, Jiang LL, et al. UHPLC/Q-TOFMS-based plasma metabolomics of polycystic ovary syndrome patients with and without insulin resistance. J Pharm Biomed Anal [Internet]. 2016;121:141–50. Available from: 10.1016/j.jpba.2016.01.025

21. Chiang JYL. Bile acid metabolism and signaling in liver disease and therapy. Liver Res [Internet]. 2017;1(1):3–9. Available from: 10.1016/j.livres.2017.05.001

22. Thomas C, Pellicciari R, Pruzanski M, Auwerx J, Schoonjans K. Targeting bile-acid signalling for metabolic diseases. Nat Rev Drug Discov. 2008;7(8):678–93.

23. Fiorucci S, Biagioli M, Zampella A, Distrutti E. Bile acids activated receptors regulate innate immunity. Front Immunol. 2018;9(AUG):1–17.

24. He C, Shan Y, Song W. Targeting gut microbiota as a possible therapy for diabetes. Nutr Res [Internet]. 2015;35(5):361–7. Available from: 10.1016/j.nutres.2015.03.002

25. Haus JM, Kashyap SR, Kasumov T, Zhang R, Kelly KR, Defronzo RA, et al. Plasma ceramides are elevated in obese subjects with type 2 diabetes and correlate with the severity of insulin resistance. Diabetes. 2009;58(2):337–43.

26. Chaurasia B, Summers SA. Ceramides - Lipotoxic Inducers of Metabolic Disorders. Trends Endocrinol Metab [Internet]. 2015;26(10):538–50. Available from: 10.1016/j.tem.2015.07.006

27. Li J, Xie LM, Song JL, Yau LF, Mi JN, Zhang CR, et al. Alterations of Sphingolipid Metabolism in Different Types of Polycystic Ovary Syndrome. Sci Rep. 2019;9(1):1– 11.

28. Mannerås-Holm L, Leonhardt H, Kullberg J, Jennische E, Odén A, Holm G, et al. Adipose tissue has aberrant morphology and function in PCOS: Enlarged adipocytes and low serum adiponectin, but not circulating sex steroids, are strongly associated with insulin resistance. J Clin Endocrinol Metab. 2011;96(2):304–11.

29. Ożegowska K, Plewa S, Mantaj U, Pawelczyk L, Matysiak J. Serum metabolomics in pcos women with different body mass index. J Clin Med. 2021;10(13).

30. Zeber-Lubecka N, Ciebiera M, Hennig EE. Polycystic Ovary Syndrome and Oxidative Stress—From Bench to Bedside. Int J Mol Sci. 2023;24(18).

31. Shukla P, Mukherjee S. Mitochondrial dysfunction: An emerging link in the pathophysiology of polycystic ovary syndrome. Mitochondrion [Internet]. 2020;52(February):24–39. Available from: 10.1016/j.mito.2020.02.006

32. Zhang J, Lu L, Tian X, Wang K, Xie G, Li H, et al. Lipidomics Revealed Aberrant Lipid Metabolism Caused by Inflammation in Cardiac Tissue in the Early Stage of Systemic Lupus Erythematosus in a Murine Model. Metabolites. 2022;12(5).

33. Zhang J, Chen L, Zheng J, Zeng T, Li H, Xiao H, et al. The protective effect of resveratrol on islet insulin secretion and morphology in mice on a high-fat diet. Diabetes Res Clin Pract [Internet]. 2012;97(3):474–82. Available from: 10.1016/j.diabres.2012.02.029

34. Siemers KM, Klein AK, Baack ML. Mitochondrial Dysfunction in PCOS: Insights into Reproductive Organ Pathophysiology. Int J Mol Sci. 2023;24(17).

35. Rajska A, Buszewska-Forajta M, Rachoń D, Markuszewski MJ. Metabolomic insight into polycystic ovary syndrome—An overview. Int J Mol Sci. 2020;21(14):1–21.

