## Supplemental Table 1 for "Metabolomic Profiling of Serum Biomarkers in Women with Polycystic Ovary Syndrome: Insights from an Untargeted Approach"

**Supplementary Table 1** Serum Metabolites found in PCOS and Control women.

| **Name of metabolites** | **Parent ion (m/z)** | **Ion mode** | **Class of compound** | **Fold Change (Control/PCOS)** | **log2 (FC)** | **Control vs PCOS** |
|---|---|---|---|---|---|---|
| PI (12:0/4:0) | 585.44 | Negative | Glycerophospholipid | 0.009 | -6.8 | ↓ |
| 3-hydroxy-eicosanoic acid | 327.33 | Negative | Fatty acyls | 60.98 | 5.93 | ↑ |
| Asn-Tyr | 294.11 | Negative | Dipeptides | 41.31 | 5.369 | ↑ |
| Tauro chenodeoxycholic acid | 522.02 | Positive | Primary bile acids | 30.42 | 4.927 | ↑ |
| Ile-Ser | 217.19 | Negative | Dipeptides | 28.58 | 4.837 | ↑ |
| Iodoform | 394.71 | Positive | Alkyl halides | 16.45 | 4.04 | ↑ |
| 3-(3,5-Diiodo-4-hydroxyphenyl) pyruvate | 430.78 | Positive | Carbohydrate derivatives | 15.78 | 3.98 | ↑ |
| PG (18:3(6Z,9Z,12Z)/19:0) | 785.34 | Negative | Glycerophospholipid | 13 | 3.7 | ↑ |
| 3'-Deoxy-cytidine-5'-triphosphate | 487.98 | Negative | Pentose phosphate | 0.093 | -3.43 | ↓ |
| 6-Methoxykaempferol 3-(6"-acetylglucoside) | 520.98 | Positive | Flavonoids | 0.11 | -3.18 | ↓ |
| Dihydrovitamin K1 | 453.42 | Positive | Prenol lipids | 8.241 | 3.043 | ↑ |
| Cer(d17:1/16:0) | 524.53 | Positive | Sphingolipids | 0.132 | -2.92 | ↓ |
| 16-methyl oleic acid | 295.17 | Negative | Fatty acid methyl ester | 6.859 | 2.778 | ↑ |

| | | | | | | |
|---|---|---|---|---|---|---|
| Bilirubin glucuronide | 783.04 | Positive | Conjugated bilirubin | 0.147 | -2.77 | ↓ |
| N-Palmitoyl histidine | 437.98 | Negative | Acylated amino acid | 6.158 | 2.622 | ↑ |
| O-(17-Carboxyheptadecanoyl) carnitine | 399.28 | Positive | Fatty Acyls | 0.166 | -2.59 | ↓ |
| TG (20:0/12:0/α-21:0) | 877.78 | Positive | Glycerolipids | 5.638 | 2.495 | ↑ |
| Taurolithocholic acid | 482.48 | Negative | Secondary bile cids | 5.54 | 2.47 | ↑ |
| Cer(d14:0/27:0) | 636.57 | Negative | Ceramide | 5.254 | 2.393 | ↑ |
| PE-NMe2(24:0/22:0) | 678.55 | Positive | Glycerophospholipid | 5.157 | 2.367 | ↑ |
| PE (18:1(9Z)-O(12,13)/24:1(15Z)) | 842.67 | Positive | Glycerophospholipid | 0.203 | -2.3 | ↓ |
| DG (22:5(4Z,7Z,10Z,13Z,19Z)-O(16,17)/0:0/i-21:0) | 643.48 | Positive | Glycerolipids | 4.66 | 2.22 | ↑ |
| FAD | 786.32 | Positive | Flavin nucleotides | 4.415 | 2.142 | ↑ |
| Ethylene glycol distearate | 595.61 | Positive | Fatty acyls | 4.304 | 2.106 | ↑ |
| 2-oxo-eicosanoic acid | 325.27 | Negative | Fatty acyls | 4.244 | 2.085 | ↑ |
| Trp-Lys-Tyr-Met-Val-D-Met | 877.36 | Negative | Peptides | 0.249 | -2.01 | ↓ |
| PC (18:1(12Z)-2OH (9,10)/18:0) | 820.64 | Positive | Glycerophospholipid | 3.819 | 1.933 | ↑ |
| PE (24:0/24:0) | 916.78 | Positive | Glycerophospholipid | 0.282 | -1.83 | ↓ |
| Thiamin Diphosphate | 446.97 | Positive | Vitamin cofactor | 3.129 | 1.646 | ↑ |

| | | | | | | |
|---|---|---|---|---|---|---|
| Phosphatidylinositol-3,4,5-trisphosphate, | 642.99 | Negative | Glycerophospholipid | 0.338 | -1.57 | ↓ |
| Fatty acyls | 450.06 | Negative | Lipids | 2.929 | 1.55 | ↑ |
| PA (20:5(6E,8Z,11Z, 14ZC265:C273,17Z)-OH(5)/i-14:0) | 683.44 | Positive | saturated fatty acid | 0.343 | -1.54 | ↓ |
| PA(P-20:0/18:1(9Z)) | 713.6 | Negative | Glycerophospholipid | 0.375 | -1.42 | ↓ |
| 25-Hydroxyvitamin D3- bromoacetate | 521.08 | Positive | Vitamin D3 derivative | 0.378 | -1.4 | ↓ |
| 5,6-dihydroxy-NADP | 780.01 | Positive | Purine nucleotides | 0.394 | -1.34 | ↓ |
| N-Acetylsphinganine | 344.36 | Positive | Sphingolipids | 2.355 | 1.236 | ↑ |
| NA-Putrescine 22:1 (11Z) | 409.49 | Positive | Acylated polyamine | 0.445 | -1.17 | ↓ |
| 6-Deoxocastasterone | 451.38 | Positive | Steroids and steroid derivatives | 0.445 | -1.17 | ↓ |
| Citrusin I | 704.42 | Positive | Carboxylic acids & derivatives | 2.152 | 1.106 | ↑ |
| Glu-His-Val | 384.18 | Positive | Carboxylic acids & derivatives | 0.491 | -1.03 | ↓ |
| PS (24:0/24:0) | 960.79 | Positive | Glycerophospholipid | 2.018 | 1.013 | ↑ |
| NA-Putrescine 17:0 | 339.37 | Negative | Fatty acyls | 0.501 | -1 | ⊗ |
| PI(O-16:0/0:0) | 557.49 | Negative | Glycerophospholipid | 1.994 | 0.995 | ⊗ |

| | | | | | | |
|---|---|---|---|---|---|---|
| PI (13:0/4:0) | 601.24 | Positive | Glycerophospholipid | 1.98 | 0.986 | ✖ |
| Kaempferol 3-O-sophoroside 7-O-glucuronide | 787.15 | Positive | Flavannoids | 0.512 | -0.97 | ✖ |
| Ile-Ser-Trp | 405.24 | Positive | Carboxylic acids and derivatives | 1.789 | 0.84 | ✖ |
| PA (12:0/13:0) | 549.22 | Negative | Glycerophospholipid | 1.74 | 0.799 | ✖ |
| PG(P-20:0/15:0) | 749.56 | Positive | Glycerolipids | 0.578 | -0.79 | ✖ |
| 2-hydroxy-tridecanoic acid | 229.22 | Negative | Fatty acids and conjugates | 1.726 | 0.787 | ✖ |
| PC(O-32:1) | 738.56 | Negative | Glycerophospholipid | 0.602 | -0.73 | ✖ |
| Octadec-5-enoic acid | 283.26 | Positive | Fatty Acyls | 1.617 | 0.694 | ✖ |
| PG (12:0/16:0) | 667.39 | Positive | Glycerophospholipid | 0.63 | -0.67 | ✖ |
| PG (24:0/ 2:0) | 637.09 | Negative | Glycerophospholipid | 0.644 | -0.63 | ✖ |
| N-Acetyl-3-methylhistidine | 212.08 | Positive | Carboxylic acids and derivatives | 0.647 | -0.63 | ✖ |
| PC (6 keto-PGF1alpha/P-18:0) | 860.61 | Positive | Glycerophospholipid | 0.648 | -0.63 | ✖ |
| 3-Hydroxyphloretin 2'-O-xylosyl-glucoside | 583.15 | Negative | Flavonoids | 0.655 | -0.61 | ✖ |
| DG (14:0/20:3(5Z,8Z,11Z)/0:0) | 589.52 | Negative | Glycerophospholipid | 1.524 | 0.608 | ✖ |
| 6-chlorotryptophan | 239.29 | Positive | Amino acid derivative | 0.682 | -0.55 | ✖ |
| GlcCer (d14:1/18:1) | 668.5 | Negative | Ceramides | 1.424 | 0.51 | ✖ |

| | | | | | | |
|---|---|---|---|---|---|---|
| DG(i-17:0/i-12:0/0:0) | 527.44 | Negative | Glycerolipids | 0.703 | -0.51 | ✖ |
| Lysophosphatidic acid | 437.53 | Positive | Glycerophospholipid | 0.713 | -0.49 | ✖ |
| 2,3-dimethyl octane | 141.2 | Negative | Fatty acyls | 0.715 | -0.48 | ✖ |
| Gln-Gln-Tyr | 436.14 | Negative | Carboxylic acids and derivatives | 0.718 | -0.48 | ✖ |
| Tetralone-4-O-Beta-D-Glucopyranoside | 323.09 | Negative | Glucosides | 1.387 | 0.472 | ✖ |
| DG (20:4(5Z,7E,11Z,14Z)-OH(9)/C2:C480:0/i-14:0), | 605.49 | Positive | Glycerolipids | 0.722 | -0.47 | ✖ |
| NA-DOPA 20:2(11Z,14Z) | 486.35 | Negative | Fatty acyls | 1.38 | 0.465 | ✖ |
| erythro-8,10-Pentatriacontanediol | 525.6 | Positive | Fatty acyls | 0.741 | -0.43 | ✖ |
| Cer(t18:0/20:3(8Z,11Z,14Z) | 638.53 | Positive | sphingosine | 0.756 | -0.4 | ✖ |
| Guanosine diphosphate adenosine | 691.15 | Negative | Nucleosides | 1.322 | 0.402 | ✖ |
| TG (16:0/20:4(5Z,8Z,11Z,14Z)/20:4(8Z,11Z,14Z,17Z)) | 903.75 | Positive | Glycerolipids | 0.762 | -0.39 | ✖ |
| GalCer (d18:2/23:0) | 794.55 | Negative | Ceramide | 1.304 | 0.383 | ✖ |
| Heme | 614.98 | Negative | Metalloporphyrin | 0.773 | -0.37 | ✖ |
| DG (22:5(4Z,7Z,10Z,13Z,19Z)-O (16,17)/0:C423:C4340/i-21:0) | 727.63 | Positive | Glycerolipids | 0.785 | -0.35 | ✖ |
| PA(i-24:0/a-25:0) | 887.73 | Positive | Glycerophospholipid | 0.801 | -0.32 | ✖ |
| TG (16:0/8:0/8:0) | 581.49 | Negative | Glycerolipids | 1.235 | 0.305 | ✖ |

| | | | | | | |
|---|---|---|---|---|---|---|
| Cer(d18:1/18:0) | 588.58 | Negative | Ceramide | 0.819 | -0.29 | ✖ |
| PI (16:0/2:0) | 613.1 | Negative | Glycerophospholipid | 0.822 | -0.28 | ✖ |
| Ile-Ser-Tyr | 382.2 | Positive | Carboxylic acids and derivatives | 0.836 | -0.26 | ✖ |
| TG (15:0/20:3(5Z,8Z,11Z)/18:0) | 871.77 | Positive | Glycerolipids | 0.844 | -0.24 | ✖ |
| Androstane | 261.25 | Positive | Steroids and steroid derivatives | 0.849 | -0.24 | ✖ |
| D-ribose 1-phosphate | 228.92 | Negative | Carbohydrate conjugate | 1.173 | 0.23 | ✖ |
| 3-ADP-2-phosphoglyceric acid | 594 | Negative | Purine nucleotides | 0.871 | -0.2 | ✖ |
| PE (12:0/13:0) | 592.29 | Negative | Glycerolipids | 1.148 | 0.199 | ✖ |
| N-acetyl serotonin glucuronide | 393.01 | Negative | Glycosides | 1.132 | 0.179 | ✖ |
| SM (d18:2/21:0) | 771.69 | Positive | Glycerolipids | 0.893 | -0.16 | ✖ |
| PS (O-14:0/20:0) | 750.64 | Positive | Lipids | 0.895 | -0.16 | ✖ |
| DG (PGD1/0:0/a-25:0) | 793.64 | Positive | Glycerolipids | 0.895 | -0.16 | ✖ |
| Spermine | 203.27 | Positive | Aliphatic peptide | 0.895 | -0.16 | ✖ |
| UDP-N-acetyl D-galactosamine 4,6-bissulfate | 785.05 | Positive | Sulfated nucleotide sugar | 0.898 | -0.15 | ✖ |
| FAHFA 13:0/3O-16:0 | 469.42 | Positive | Fatty esters | 0.899 | -0.15 | ✖ |
| DG (22:5(4Z,7Z,10Z,13Z,19Z)- | 727.63 | Positive | Glycerolipids | 0.899 | -0.15 | ✖ |

| | | | | | | |
|---|---|---|---|---|---|---|
| O(16,17)/0C146:C171:0/i-15:0) | | | | | | |
| DG (20:5(7Z,9Z,11E,13E,17Z)-3OH (5,6,15)/0:0/10:0) | 701.61 | Positive | Glycerolipids | 1.111 | 0.151 | ⊗ |
| 5-Bromo-DL-tryptophan | 280.93 | Negative | Indoles and derivatives | 0.902 | -0.15 | ⊗ |
| omega-linoleoyloxy-Cer(d18:1/36:0) | 1095.08 | Negative | Ceramides | 0.902 | -0.15 | ⊗ |
| PI-Cer(d14:1(4E)/21:0) | 792.49 | Negative | Sphingolipids | 0.903 | -0.15 | ⊗ |
| Phosphoceramide | 796.52 | Negative | Ceramides | 1.107 | 0.146 | ⊗ |
| 4-amino-2-methyl-5-diphosphomethylpyrimidine | 294.95 | Negative | Pyridines and derivatives | 0.909 | -0.14 | ⊗ |
| PE (12:0/17:2(9Z,12Z)) | 644.25 | Negative | Glycerophospholipid | 0.91 | -0.14 | ⊗ |
| PI (O-14:0/10:0) | 683.49 | Negative | Glycerophospholipid | 0.913 | -0.13 | ⊗ |
| UDP-N-acetylglucosamine | 606.14 | Negative | Pyrimidine nucleotides | 0.913 | -0.13 | ⊗ |
| Vitamin D3 butyrate | 453.44 | Negative | Sterol lipid | 0.913 | -0.13 | ⊗ |
| 5-Diphosphoinositol pentakisphosphate | 740.83 | Positive | Organooxygen compounds | 0.914 | -0.13 | ⊗ |
| Cinnamoyl-CoA | 915.17 | Positive | Acyl-CoA | 0.914 | -0.13 | ⊗ |
| FAHFA (18:0/8-O-18:0) | 584.59 | Negative | Fatty esters | 0.914 | -0.13 | ⊗ |
| TG (14:1(9Z)/16:0/18:1(9Z)) | 803.74 | Positive | Glycerolipids | 0.915 | -0.13 | ⊗ |

| | | | | | | |
|---|---|---|---|---|---|---|
| 2-Aminoadenosine | 281.1 | Negative | Purine nucleosides | 0.915 | -0.13 | ⊗ |
| DG (20:5(6E,8Z,11Z,14Z,17Z) | 701.61 | Positive | Glycerolipids | 0.918 | -0.12 | ⊗ |
| NAD | 664.13 | Positive | Nucleosides | 0.918 | -0.12 | ⊗ |
| Cer(t18:0/20:4(7E,9E,11Z,13E)-3OH(5S,6R,15S)) | 652.56 | Positive | Sphingolipids | 1.083 | 0.115 | ⊗ |
| 4-Hydroxy-17beta-estradiol-2-S-glutathione | 594.28 | Positive | Steroids and steroid derivatives | 0.936 | -0.1 | ⊗ |
| Tresperimus | 388.29 | Positive | Carboxylic acids & derivatives | 1.065 | 0.091 | ⊗ |
| Cer (d18:1/5-iso PGF2VI) | 608.5 | Positive | Sphingolipids | 1.064 | 0.09 | ⊗ |
| Cer (d17:1/18:1(9Z)-O (12,13)) | 564.45 | Positive | Sphingolipids | 1.061 | 0.085 | ⊗ |
| 1"-O-Beta-D-Glucopyranosylformoside | 685.19 | Negative | Prenol lipids | 1.058 | 0.081 | ⊗ |
| 16:3 Glc-Cholesterol | 781.1 | Positive | Glucosylated cholesterol | 1.046 | 0.065 | ⊗ |
| Pomolic Acid-28-O-Beta-D-Glucopyranosyl Ester | 633.23 | Negative | Triterpenoids | 1.046 | 0.065 | ⊗ |
| Cer (d18:2/ 18:1) | 600.96 | Positive | Ceramide | 1.046 | 0.064 | ⊗ |
| 9-Octadecenoic acid | 282.29 | Positive | Fatty acyls | 1.046 | 0.064 | ⊗ |
| SM (d19:0/5-iso PGF2VI) | 789.54 | Positive | Ceramide | 1.042 | 0.06 | ⊗ |
| PG (O-14:0/ 38:0) | 971.72 | Positive | Glycerophospholipid | 1.042 | 0.06 | ⊗ |
| 4,5-Dibromo-1-ethylpyrrole-2-carboxylic acid | 293.92 | Negative | Pyrroles | 1.042 | 0.06 | ⊗ |

| | | | | | | |
|---|---|---|---|---|---|---|
| Phosphoaminophosphonic acid-guanylate ester | 523.02 | Positive | Purine nucleotides | 1.042 | 0.059 | ✖ |
| TG (14:0/22:2(13Z,16Z)/20:3(5Z,8Z,11Z)) | 909.77 | Positive | Glycerolipids | 1.04 | 0.057 | ✖ |
| TG (18:2(9Z,12Z)/24:1(15Z)/O-18:0) | 955.92 | Positive | Glycerolipids | 1.04 | 0.057 | ✖ |
| PG (22:0/2:0) | 609.21 | Negative | Glycerophospholipid | 1.038 | 0.053 | ✖ |
| Inositol 1,3,4,5,6-pentakisphosphate | 578.9 | Negative | Inositol phosphate | 1.036 | 0.051 | ✖ |
| His-Val-Val | 352.24 | Negative | Carboxylic acids & derivatives | 1.036 | 0.051 | ✖ |
| N-oleoyl asparagine | 395.04 | Negative | N-acylamide | 1.036 | 0.051 | ✖ |
| PC (20:4(8Z,11Z,14Z,17Z)-2OH(5S,6R)/14:1(9Z)) | 782.5 | Negative | Glycerolipids | 0.966 | -0.05 | ✖ |
| GDP-guluronate | 618.04 | Positive | Purine nucleotides | 1.034 | 0.049 | ✖ |
| Gymnodimine | 508.33 | Positive | Pyridines and derivatives | 1.031 | 0.044 | ✖ |
| PE-NMe2(15:0/14:0) | 678.55 | Negative | Glycerophospholipid | 1.029 | 0.041 | ✖ |
| PI (P-16:0/14:0) | 767.6 | Positive | Glycerolipids | 1.028 | 0.04 | ✖ |
| UDP N-acetyl-D-galactosamin 4-sulfate | 705.18 | Positive | Pyrimidine nucleotide sugars | 1.024 | 0.034 | ✖ |
| PA(P-16:0/14:0) | 603.25 | Negative | Glycerolipids | 1.024 | 0.034 | ✖ |
| 17-Beta-Estradiol-3,17-beta-sulfate | 433.04 | Positive | steroid conjugates | 0.99 | -0.01 | ✖ |
| GDP-mannuronate | 617.01 | Positive | Purine nucleotides | 1.003 | 0.004 | ✖ |

| | | | | | | |
|---|---|---|---|---|---|---|
| Coenzyme A | 766.07 | Negative | Coenzyme | 1.003 | 0.004 | ⊗ |
| ↓ Significant downregulation | ↑ Significant upregulation | | | ⊗ Unsignificant | | |
