## Supplemental Table 2 for "Metabolomic Profiling of Serum Biomarkers in Women with Polycystic Ovary Syndrome: Insights from an Untargeted Approach"

**Supplementary Table 2**. T test analysis of identified serum metabolites of participants.

| Name of metabolites | t. stat | p-value | p (-log10) | FDR |
|---|---|---|---|---|
| PG (18:3(6Z,9Z,12Z)/19:0) | 25.765 | 2.11E-51 | 50.675 | 2.92E-49 |
| Taurolithocholic acid | 14.465 | 2.60E-28 | 27.585 | 1.79E-26 |
| 3-hydroxy-eicosanoic acid | 11.565 | 2.18E-21 | 20.661 | 1.00E-19 |
| Cer(d14:0/27:0) | 7.8503 | 1.74E-12 | 11.76 | 5.99E-11 |
| PI (12:0/4:0) | -7.4723 | 1.27E-11 | 10.895 | 3.51E-10 |
| PA (12:0/13:0) | 7.2065 | 5.06E-11 | 10.296 | 1.16E-09 |
| Dihydrovitamin K1 | 6.5716 | 1.26E-09 | 8.898 | 2.49E-08 |
| 16-methyl oleic acid | 6.2766 | 5.40E-09 | 8.2677 | 9.31E-08 |
| Iodoform | 5.4003 | 3.30E-07 | 6.4817 | 5.06E-06 |
| 3'-Deoxy-cytidine-5'-triphosphate | -5.2183 | 7.43E-07 | 6.129 | 1.03E-05 |
| FAD | 4.8893 | 3.09E-06 | 5.5095 | 3.72E-05 |
| Cer(d17:1/16:0) | -4.879 | 3.23E-06 | 5.4907 | 3.72E-05 |
| Tauro chenodeoxycholic acid | 4.6865 | 7.24E-06 | 5.14 | 7.69E-05 |
| PA (20:5(6E,8Z,11Z,14ZC265:C273,17Z)-OH(5)/i-14:0) | -4.5467 | 1.29E-05 | 4.8909 | 0.0001 |
| 6-Methoxykaempferol 3-(6''-acetylglucoside) | -4.4504 | 1.90E-05 | 4.7222 | 0.0001 |
| Phosphatidylinositol-3,4,5-trisphosphate, | -4.2961 | 3.49E-05 | 4.4566 | 0.0003 |
| PE (24:0/24:0) | -4.1581 | 5.97E-05 | 4.2242 | 0.0004 |
| PE (18:1(9Z)-O(12,13)/24:1(15Z)) | -4.0897 | 7.75E-05 | 4.1108 | 0.0005 |
| PE-NMe2(24:0/22:0) | 3.9855 | 0.00011 | 3.9408 | 0.0008 |
| TG (20:0/12:0/a-21:0) | 3.9161 | 0.00014 | 3.8291 | 0.0010 |
| Asn-Tyr | 3.8279 | 0.00020 | 3.689 | 0.0013 |

| | | | | |
|---|---|---|---|---|
| O-(17-Carboxyheptadecanoyl) carnitine | -3.8212 | 0.00020 | 3.6785 | 0.0013 |
| 25-Hydroxyvitamin D3-bromo acetate | -3.7549 | 0.00026 | 3.5748 | 0.0015 |
| 6-Deoxocastasterone | -3.7502 | 0.00027 | 3.5674 | 0.0015 |
| Trp-Lys-Tyr-Met-Val-D-Met | -3.7236 | 0.00029 | 3.5263 | 0.0016 |
| 2-oxo-eicosanoic acid | 3.7184 | 0.00030 | 3.5182 | 0.0016 |
| NA-Putrescine 22:1 (11Z) | -3.3299 | 0.00114 | 2.9404 | 0.0057 |
| DG (22:5(4Z,7Z,10Z,13Z,19Z)-O(16,17)/0:0/i-21:0) | 3.3267 | 0.00115 | 2.9359 | 0.0057 |
| 2,2,3,3,4,4,5,5,6,6,7,7,8,8,9,9-Hexadecafluorononanoic acid | 3.1643 | 0.00195 | 2.7079 | 0.0093 |
| 5,6-dihydroxy-NADP | -3.1209 | 0.00224 | 2.6484 | 0.0103 |
| NA-Putrescine 17:0 | -3.0669 | 0.00266 | 2.575 | 0.0118 |
| PC(O-32:1) | -3.0357 | 0.00292 | 2.5332 | 0.0126 |
| 3-(3,5-Diiodo-4-hydroxyphenyl) pyruvate | 2.9699 | 0.00358 | 2.4458 | 0.0149 |
| Glu-His-Val | -2.948 | 0.00382 | 2.417 | 0.0155 |
| 3-Hydroxyphloretin 2'-O-xylosyl-glucoside | -2.9263 | 0.00408 | 2.3886 | 0.0160 |
| PA(P-20:0/18:1(9Z)) | -2.9188 | 0.00417 | 2.3789 | 0.0160 |
| Bilirubin glucuronide | -2.7866 | 0.00617 | 2.2096 | 0.0226 |
| 2,3-dimethyloctane | -2.7831 | 0.00623 | 2.2052 | 0.0226 |
| PG (24:0/ 2:0) | -2.7443 | 0.00697 | 2.1566 | 0.0246 |
| Ile-Ser | 2.7252 | 0.00736 | 2.1328 | 0.0248 |
| PS (24:0/24:0) | 2.7238 | 0.00739 | 2.1311 | 0.0248 |
| Kaempferol 3-O-sophoroside 7-O-glucuronide | -2.6864 | 0.00822 | 2.0851 | 0.0270 |
| Gln-Gln-Tyr | -2.5689 | 0.01139 | 1.9432 | 0.0365 |

| | | | | |
|---|---|---|---|---|
| Cer(t18:0/20:3(8Z,11Z,14Z) | -2.4764 | 0.01463 | 1.8348 | 0.0446 |
| erythro-8,10-Pentatriacontanediol | -2.4715 | 0.014822 | 1.8291 | 0.0446 |
| DG(i-17:0/i-12:0/0:0) | -2.4703 | 0.014869 | 1.8277 | 0.0446 |
| 4-Methyltriacontane | -2.439 | 0.016157 | 1.7916 | 0.0474 |
| Cer(d18:1/18:0) | -2.4142 | 0.017244 | 1.7634 | 0.0486 |
| Heme | -2.4085 | 0.017504 | 1.7569 | 0.0486 |
| PG (12:0/16:0) | -2.406 | 0.017615 | 1.7541 | 0.0486 |
